# A bifunctional CPD/(6-4)- photolyase from the cyanobacteria *Synechococcus* sp. PCC 7335 constitutes an UV-B inducible operon: new insights into photolyases phylogenetic evolution

**DOI:** 10.1101/2022.06.29.498140

**Authors:** Fernández María Belén, Latorre Lucas, Correa-Aragunde Natalia, Raúl Cassia

**Affiliations:** Instituto de Investigaciones Biológicas- Facultad de Ciencias Exactas y Naturales, Universidad Nacional de Mar Del Plata- Consejo Nacional de Investigaciones Científicas y Técnicas, CC1245 7600, Mar Del Plata, Buenos Aires, Argentina

**Keywords:** Bifunctional CPD/(6-4)- photolyase, Cryptochrome/ photolyase family, *Synechococcus* sp. PCC 7335, UV-B, Photolyases operon, Cyanobacteria

## Abstract

Photosynthetic organisms are continuously exposed to solar ultraviolet radiation-B (UV-B) because of their autotrophic lifestyle. UV-B provokes DNA damages, as cyclobutane pyrimidine dimers (CPD) or pyrimidine (6-4) pyrimidone photoproducts (6-4 PPs). The cryptochrome/ photolyase family (CPF) comprises flavoproteins that are able to bind damaged or undamaged DNA. Photolyases (PHRs) are enzymes that repair either CPDs or 6-4 PPs. A natural bifunctional CPD/(6-4)- PHR (PhrSph98) was recently isolated from the UV-resistant bacteria *Sphingomonas* UV9. In this work, phylogenetic studies of bifunctional CPD/(6-4)- photolyases and its evolutionary relationship with other CPF members were performed. Amino acids involved in electron transfer and binding to FAD cofactor and DNA lesions were conserved in proteins from proteobacteria, planctomycete, bacteroidete, acidobacteria and cyanobacteria clades. Genome analysis revealed that the cyanobacteria *Synechococcus* sp. PCC 7335 encodes a two-gene assembly operon coding for a PHR and a bifunctional CPD/(6-4)- PHR. Operon structure was validated by RT-qPCR analysis and the polycistronic transcript accumulated after 15 minutes of UV-B irradiation. Conservation of structure and evolution is discussed. This is the first report of an UV-B inducible PHRs operon which encodes a CPD/(6-4)- photolyase with putative bifunctional role in the repair of CPDs and 6-4 PPs damages in oxygenic photosynthetic organisms.

## 1 Introduction

Ultraviolet-B (UV-B) is the solar electromagnetic radiation with wavelengths between 280-315 nm. Although most of this radiation is absorbed by the stratospheric ozone layer it affects all living organisms. Around 0.3% of sunlight energy at sea level corresponds to UV-B and it can damage aquatic organisms triggering a decrease of the ecosystem productivity (Häder et al., 2011; Banaś et al., 2020). UV-B is perceived by the UV-B response locus 8 (UVR8) photoreceptor present in photosynthetic organisms ranging from green algae to higher plants (Fernández et al., 2016). Although no defined photoreception systems have been described for other photosynthetic microorganism (red and brown algae and cyanobacteria), they have developed several mechanisms of protection against UV-B damage (Montgomery, 2007; Sinha and Häder, 2008; Rastogi et al., 2014).

Cyanobacteria are the most primitive group of Gram-negative bacteria, and the first prokaryotes that perform oxygenic photosynthesis. Cyanobacteria are ubiquitous and occupy diverse ecological niches, adapting to various extreme environments, such as high or low temperatures, highly acidic or basic pH, high salt concentrations, desiccation and UV-B (Wada et al., 2013). As photosynthetic organisms, they depend of solar energy and have to cope with harmful UV-B. Solar UV-B affects the DNA and protein structures, photosynthesis, ribulose 1,5-bisphosphate carboxylase/oxygenase (RuBisCO) activity, N_2_ fixation, cellular morphology, growth, survival, pigmentation and buoyancy (Tyagi et al., 1992; Rastogi et al., 2014; Kumar et al., 2020; Vega et al., 2020). Thus, cyanobacteria developed several UV protection mechanisms that include UV absorbing and screening compounds as scytonemin and mycosporine-like amino acids (MAAs), antioxidant protection, apoptosis, migration, mat formation and DNA reparation to recover UV induced DNA damages (Sinha and Häder, 2008; Moon et al., 2012; Rastogi et al., 2014).

Solar UV radiation can induce two types of pyrimidine dimers in the double helix DNA, being the predominant cyclobutane pyrimidine dimers (75%) (CPDs) and to a lesser extent pyrimidine (6-4) pyrimidone photoproducts (6-4 PPs) (Banaś et al., 2020; Vechtomova et al., 2020). Photoreactivation is a blue/ UV-A light-dependent mechanism used to specifically repair CPD or 6-4 PPs damages by photolyases (PHRs) (Pathak et al., 2019). These enzymes are a class of flavoproteins found in all organisms excluding placental mammals, who lost all genes encoding functional photolyases in the course of evolution (Xu et al.; Banaś et al., 2020). During repair, flavin adenine dinucleotide (FAD) cofactor is fully reduced via Trp or Tyr surface transfer of electrons in a process named as photoreduction (7). Then, a rapid electron transfer from the excited fully reduced FAD chromophore to the DNA lesion triggers both kind of repair. The 6-4 PPs repair also requires proton transfer which is a limiting step resulting in a much lower reaction efficiency (1). A bifunctional photolyase called as PhrSph98 was recently characterized in the Antarctic bacterium *Sphingomonas* sp. UV9. This enzyme is able to repair both types of damages since it displays a larger catalytic pocket compared to CPDs or 6-4 PPs repairing enzymes (8).

The cryptochrome/ photolyase family (CPF) comprises proteins that have conserved the FAD binding site and switches between basal and excited states. CPF proteins bind either damaged or undamaged DNA and are classified in two groups according to their function: 1) CPD and 6-4 PPs photolyases, which repair CPD or 6-4 PPs damages respectively and 2) Cryptochromes (CRY) that regulate growth and development in plants and the circadian clock both in plants and animals (Mei and Dvornyk, 2015; Vechtomova et al., 2020).

CPD PHRs are classified, based on sequence similarity in i) class 0, which repair CPD damages in single-stranded DNA (ssDNA PHR, previously classified as CRY with Cry-DASH designation), ii) class I, present mostly in unicellular organisms; iii) class II, in unicellular and multicellular organisms; and iv) class III found only in some eubacteria (Ozturk, 2017; Zhang et al., 2017). Recently, an exhaustive phylogenetic analysis identify a new class of CPD repair enzymes, called short photolyase-like (SPL), because of the lack the N-terminal α/β domain of normal photolyases. They are similar to class I/III CPD PHRs and authors speculated that this class constitutes the real ancestor of the CPF (10). CPD PHRs III may be considered as an intermediate form between CPD PHR I and plant CRYs, as well as a protein that has retained the traits of their common ancestor (Vechtomova et al., 2020). The iron–sulfur cluster containing bacterial cryptochromes and 6-4 PPs repair photolyases (FeS – BCPs) are a newly characterized CPF members containing the amino acids necessary to bind cofactors, and four conserved cysteine residues for the coordination of an iron -sulfur cluster, an ancient feature (Scheerer et al., 2015).

Some CPFs have dual roles, like ssDNA PHR proteins. They are restricted to UV lesions and repair CPD damages in single-stranded DNA, having also blue light photoreceptor activity. These proteins may represent the link between photolyases and cryptochromes (Kiontke et al., 2020). It has been proposed that the bifunctional CPD/(6-4)- PHR from *Sphingomonas* sp. UV9, PhrSph98, is a sister group of CPD class II PHRs and that may represent a missing link in the transition from 6-4 PP to CPD PHRs (Marizcurrena et al., 2020).

It is widely accepted that photolyases are ancient DNA repair enzymes, which have evolved far before the oxygen accumulation and the ozone layer establishment in the atmosphere (Vechtomova et al., 2020). However, the evolutionary scenario was not fully elucidated yet. Prokaryotic 6-4 photolyases were suggested as the first common ancestor of photolyases. However, as CPDs are the major UV-induced DNA damage it is not convincing that the 6-4 photo repair occurred earlier than the CPD one during evolution (Xu et al., 2021). Exhaustive phylogenetic trees do not include the recently described bifunctional CPD/(6-4)- photolyase, and its occurrence among other species was not explored. Thus, the aim of this work was to analyze the distribution of CPD/(6-4)- photolyases and its evolutionary relationship with other CPF family members. We show that the cyanobacteria *Synechococcus* sp. PCC 7335 encodes a PhrSph98 homolog, in a two-gene assembly operon, which is induced by UV-B light. Evolutionary and structural aspects of this operon are discussed.

## 2 Methods

### 2.1 Defined criteria for the homology search

A protein–protein BLAST (BlastP) analysis was performed against the non-redundant protein sequence (nr) database using as template the amino acid sequence of PhrSph98 from *Sphingomonas* sp. UV9 (accession number: NCBI ANW48627), and restricting the maximum target sequences to 250. All the retrieved sequences match the criteria of E-values lower than 0.001 (being lower to 1×10^-63^) and percentage identity higher than 30% according to Pearson (2013). Subsequently, partial sequences and those sharing >85% of identity (redundant proteins) were identified and discarded using CD-HIT software (http://weizhong-lab.ucsd.edu/cdhit-web-server/cgi-bin/index.cgi?cmd=cd-hit).

### 2.2 Phylogenetic analysis and protein conserved domain search

Multiple sequence alignment and curation was performed using MAFFT (https://mafft.cbrc.jp/alignment/server/) and BMGE 1.12_1 (https://ngphylogeny.fr/tools/tool/273/form) software respectively using default settings. Subsequently, the phylogenetic tree was inferred with PhyML 3.0 (http://www.atgc-montpellier.fr/phyml/) using the automatic model selection AIC, which defined LG5+G+I+F as the best model of evolution, and 1000 bootstrap. The tree was visualized using iTOL (https://itol.embl.de/login.cgi).

A sequence alignment logo for the proteins described as homologs of PhrSph98 was created using the online available WebLogo tool (https://weblogo.berkeley.edu/logo.cgi) (Crooks et al., 2004).

Conserved domains were identified using the sequences from each target protein employing the NCBI conserved domain database (CDD)-NIH.

### 2.3 Operon prediction and sequence analysis

Operon prediction was performed using the webservers Operon mapper (https://biocomputo.ibt.unam.mx/operon_mapper/) (Taboada et al., 2018) and SoftBerry (http://www.softberry.com/berry.phtml?topic=index&group=programs&subgroup=gfindb) restricting the search to bacterial genomes.

Operon promoter and potential transcription factors binding sequences were predicted using BPROM software from SoftBerry platform (http://www.softberry.com/). Transcriptional terminator sequences were predicted using iTerm-PseKNC/predictor (http://lin-group.cn/server/iTerm-PseKNC/pre.php#) (Feng et al., 2019).

### 2.4 *Structure prediction of putative bifunctional CPD/(6-4)- PHR from* Synechococcus sp. PCC 7335

The linear amino acid sequence of PhrSph98 (ANW48627.1) was used as template in BlastP against the nr protein database restricting the search to *Synechococcus* sp PCC 7335. Tertiary structure of PhrSph98 and *Synechococcus* sp. PCC 7335 homolog were predicted using Modeller in the HHPred server (https://toolkit.tuebingen.mpg.de/tools/hhpred) (Söding et al., 2005). The structure of class II CPD-photolyase from *Methanosarcina mazei* (PDB ID: 2XRY) was the most similar to both queries, and thus selected as template. Visualization, structure overlaps and distance calculations were done using the Molecular Graphics System PyMOL.

### 2.5 Bacterial strain and culture method

The cyanobacterial strain *Synechococcus* sp. PCC 7335 used in this study was acquired from the Pasteur Culture Collection of Cyanobacteria (http://www.pasteur.fr/). *Synechococcus* sp. PCC 7335 was originally isolated from a snail shell in an intertidal zone near Puerto Peñasco, Mexico (Rippka et al., 1979). PHR genes were identified in the genome of *Synechococcus* sp. PCC 7335 (NCBI accession number NZ_DS989904.1). Cultures were grown in Erlenmeyer flasks in volumes of 150 mL containing ASNIII marine medium at 25 °C and photon flux density of 5 μmol m^-2^ sec^-1^ under a photoperiod of 12 h light (4500K LED tubes): 12 h dark. The optical density at 750 nm (OD750 nm) was measured with GeneQuant 1300 spectrophotometer to monitor cell growth.

### 2.6 UV-B treatment

*Synechococcus* sp. PCC 7335 cells were grown in at least three replicate cultures to an OD750 nm ~0.2-0.4. Aliquots of cells (~35 mL) were collected from each replicate culture, placed in Petri dishes and exposed to 3.34 μmol m^-2^ sec^-1^ of UV-B supplemented with ~5 μmol m^-2^ sec^-1^ of PAR for 15 and 30 minutes in a controlled environment chamber. The UV-B irradiance intensities were chosen according to He et al., (2021). Control samples were covered with a polycarbonate filter of 1 mm to screen UV-B. UV-B was provided by narrowband UV-B Philips TL 100W/01 lamps. The spectral irradiance was determined with an UV-B photo-radiometer (Delta ohm HD2102.1).

After treatment, cells were immediately pelleted by centrifugation at 11000 g and 4 °C and stored at - 80 °C until being processed for total RNA extraction.

### 2.7 RNA extraction and RT-qPCR analysis

RNA from *Synechococcus* sp. PCC 7335 was extracted with the RNeasy mini Kit (Qiagen) following the manufacturer’s protocol. The concentration and purity of the isolated RNA was measured with a UV spectrophotometer (NanoDrop™ One, Thermo Scientific). Total RNA was used for cDNA synthesis in a reaction containing 3μM random primer, 10 mM dNTP, 0.1 M DTT and 200 U of M-MLV reverse transcriptase (Invitrogen).

RT-qPCR was performed a 10 μL total reaction volume including 1 μL of a dilution of cDNA template, 250 nM of forward and reverse primers and 5 μL of 2x Power SYBR Green PCR Mix (Applied Biosystem). Amplified signals were monitored continuously with a Step One Plus Real-Time Thermal Cycler (Applied Biosystem). Thermocycling was performed using the following conditions: 10 min of denaturation and enzyme activation at 95 °C, followed by 40 cycles at 95 °C for 15 s and 60 °C for 1 min. Melting curve analysis was performed from 60 to 95 °C at 0.3 °C increment to verify primer specificity. Negative controls (no template and minus RT transcriptase) were included in every PCR run to verify no genomic DNA contamination. A default threshold of 1 was used. The size of qPCR products for each primer pair was verified on 2% agarose gel electrophoresis using 1x TAE buffer. Ladder 100 pb was used as molecular marker (PB-L productos Bio- lógicos). Real time data was analyzed using the StepOne™ Software v2.3 Tool (Applied Biosystem). Primers used are listed in Supplementary table 1. RNase P RNA (*RNPB*) and Phosphoenolpyruvate carboxylase (*PPC*) were used as reference genes for gene expression normalization. Results were expressed as 2^(-ΔCt)^ being ΔCt the ratio between target gene and the geometric mean of *PPC* and *RNPB* (Livak and Schmittgen, 2001).

## 3 Results

### 3.1 BLASTp analysis and amino acid conservation

In order to analyze phylogenetic positioning of the CPD/(6-4)- photolyase from *Sphingomonas* sp. UV9 (PhrSph98) we performed a BLASTp search of the NCBI non-redundant protein database. Results retrieved 250 sequences with similarities ranging from 90 to 31% and E-values from 0 to 1.10^-63^ matching the criteria of homology reported previously (Pearson, 2013). *Arabidopsis* UVR2 protein, is a class II CPD PHR that repairs CPD lesions upon UV-B irradiation and was included in the tree inference (Landry et al., 1997). MSA and phylogenetic analysis showed that the closest homologs to *Sphingomonas* sp. UV9 bifunctional CPD/(6-4)- PHR were grouped in a clade (numbered as 7, 8, 9 and 10) containing proteins with E-values of 0 and percentages of identity ranging from 61.32 to 82.04. These proteins correspond to Gram negative bacteria, including *Sphingomonas* as the most abundant (31/37), and others like *Belnapia* (1/37), *Dankookia* (1/37), *Sphingosinicella* (1/37), *Methylobacterium* (1/37), *Polymorphobacter* (1/37) and *Croceibacterium* (1/37) (Figure 1A). According to the domain profile search using the NCBI conserved domain database (CDD)- NIH, the majority of these proteins contain the PHR2 domain, except for two proteins that have a PHRB domain (WP_157177235.1 from *Sphingomonas prati* and WP_093400154.1 from *Sphingomonas jinjuensis*).

**Figure 1.**
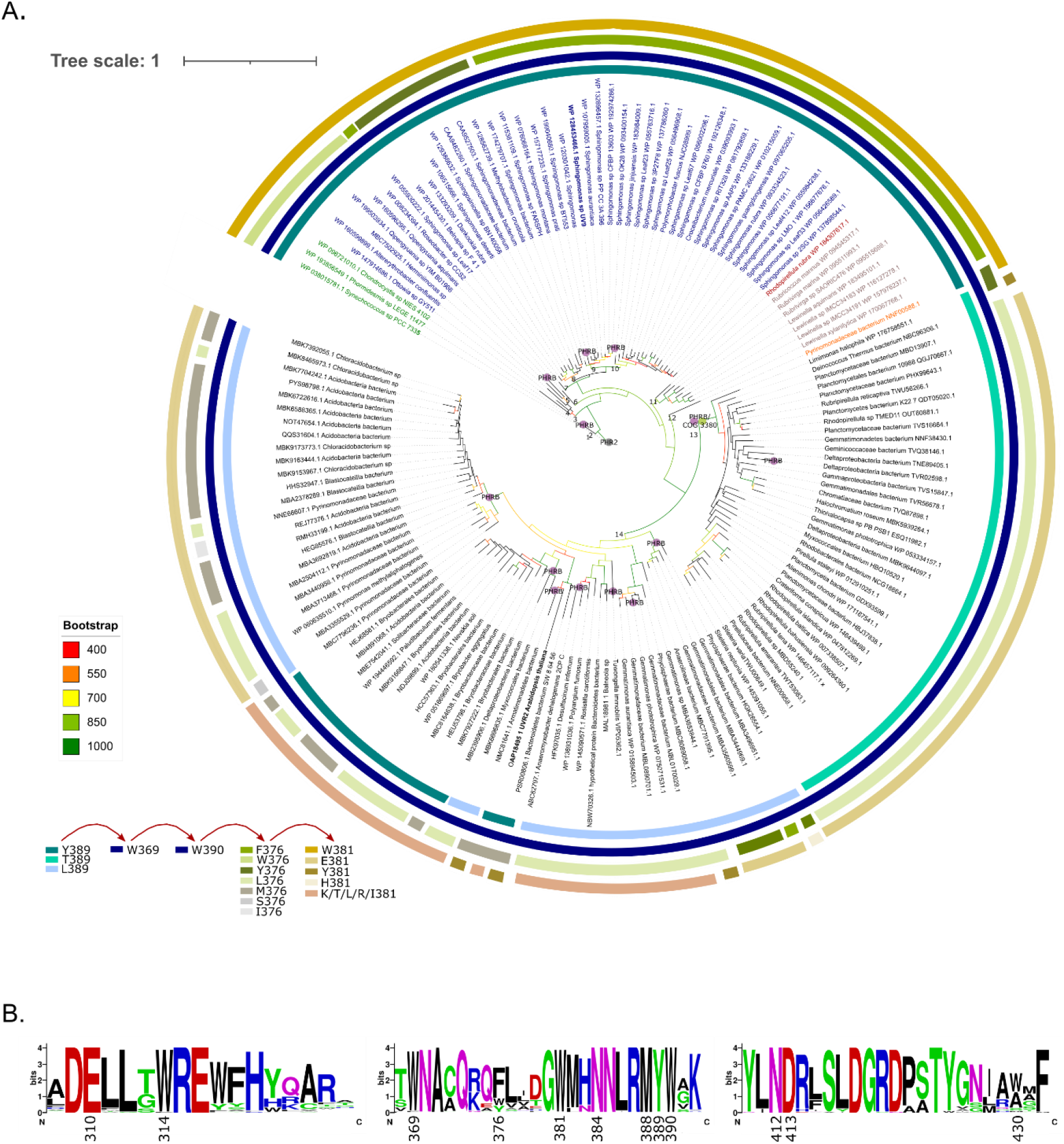
Unrooted maximum-likelihood tree of bifunctional CPD/(6-4)- PHRs. **A.** Protein sequences with similarity to PhrSph98 from *Sphingomonas* sp. UV9 were retrieved using BLASTp and the non-redundant database. CD-HIT software (Huang et al., 2010) was used to remove all sequences with identities >85% (redundant proteins). The sequences were aligned with MAFFT (http://mafft.cbrc.jp/alignment/server/). Selection of phylogenetic informative regions from the multiple sequence alignment was performed using the BMGE 1.12_1 software (Criscuolo and Gribaldo, 2010). Maximum likelihood phylogenetic tree was calculated with LG +G+ I+ F model using PHYML 3.0 (Guindon et al. 2010) and is presented as unrooted. Nonparametric bootstrapping (1000 replicates) was used to assess tree branching support. The program iTol was used to display phylogenetic trees (http://itol.embl.de/) (Letunic and Bork, 2021). Rings of different colors indicate the amino acids described in PhrSph98 to be potentially involved in electron transfer pathway: Y389- W369- W390- W376- W381. Red arrows indicate the electron flux. Protein domains PHRB, PHR2 and COG3380 are shown in each node. Text in color indicate bacteria taxonomic classification of predicted homologs of PhrSph98: green, cyanobacteria; blue, proteobacteria; red, planctomycete; brown, bacteroidete; orange, acidobacteria. The accession number of each sequence is given next to the species name. Bootstrap values are shown in colors being red the lowest (400) and green the highest (1000). Bootstrap values lower than 40% are not shown. **B.** MSA from predicted bifunctional photolyases was used to create a sequence alignment logo showing conservation of amino acids involved in electron transfer (Y389-W369-W390-F376-W381), FAD binding (W412, D413), lesion binding (W314, A430, M388) and lesion stabilization (E310, N384) using online available WebLogo tool (https://weblogo.berkeley.edu/logo.cgi) (Crooks et al., 2004). See text for additional details. The height of each letter is proportional to the frequency of the corresponding amino acid and the overall height of each stack for a position is proportional to the sequence conservation, measured in bits. N at the left and C at the right indicate amino- and carboxi-terminal regions. Numbers indicate amino acid position in PhrSph98. Polar amino acids (G,S,T,Y,C,Q,N) are shown in green, basic (K,R,H) in blue, acidic (D,E) in red and hydrophobic (A,V,L,I,P,W,F,M) amino acids are in black.

#### 3.1.1 Amino acids involved in electron transfer

*In silico* proposed model from *Sphingomonas* sp. UV9 PhrSph98 suggests that electron transfer to FAD involves Y389-W369-W390-F376-W381-FAD pathway (Marizcurrena et al., 2020). We analyzed the conservation of amino acids involved in electron transfer using a MSA. Figure 1B shows that W369 and 390 were conserved in the 149 sequences analyzed. Y389 is conserved from nodes 1 to 12, being changed for a Thr or Leu (which could not participate in the electron transfer) in the species from nodes 13 and 14. F376 was conserved in nodes 7, 9, 10 and 11 and in WP_115381109 from *Sphingomonas* sp. strain FARSPH in node 8. It was replaced for a Trp in nodes 1 to 6, and for a Tyr in nodes 8 and 12. In the other predicted proteins, there waa a change to non-aromatic amino acids in this position. Finally, W381 was conserved from node 1 to 12. Thus, aromatic amino acids important for electron transfer and DNA repair are conserved in species from nodes 1 to 12, suggesting that they constitute PhrSph98 homologs.

#### 3.1.2 FAD binding domain

In class II CPD PHRs, N403 is highly conserved and involved in FAD cofactor binding (Kiontke et al., 2011). This residue from *M. mazei* class II photolyase (template for PhrSph98 tertiary structure modeling) corresponds to N412 in PhrSph98 (Marizcurrena et al., 2020). MSA analysis showed that this residue was conserved in all the sequences predicted as homologous by phylogeny (Figure 1B). A charge compensation during FADH- photoreduction is performed by H- bonding of N403 and the surface exposed D404 in *M. mazei* CPD PHR (Kiontke et al., 2011). This residue (D413 in PhrSph98 from *Sphingomonas*) was also conserved in PhrSph98 homologs indicating that FAD binding is favored in these enzymes.

#### 3.1.3 Amino acids involved in DNA lesion binding

CPD and 6-4 PPs DNA lesions are predicted to be positioned in the binding pocket of PhrSph98 through interaction with the hydrophobic residues W314, A430 and M388 (Marizcurrena et al., 2020). W314 was conserved among all the proteins identified as PhrSph98 homologs (Figure 1B). In nodes 1 to 8 and 11 to 13 the position of A430 was replaced for a Trp, as occurs in *M. mazei* CPD photolyase II (W421) (except for WP_095511993.1, WP_095515698.1 from node 11 and TWT53083.1 from node 13). In nodes 9 and 10, A430 was conserved except for WP_157177235.1, WP_107959005.1, WP_133188229.1, WP_010215059.1, WP_056426589.1. M388 was conserved among all the protein sequences analyzed, except for protein TWT53083.1.

#### 3.1.4 Amino acids involved in lesion stabilization

Amino acids E310 and N384 from PhrSph98 are hypothesized to be involved in the stabilization of i) the CPD radical after electron transfer (by the transfer of a proton from a neutral Glu at the bottom of the active site) and ii) the anionic thymine radical after bond breakage (through hydrogen bond formation with the N3 amide and C4 carbonyl from CPD lesion) respectively (Marizcurrena et al., 2020). Both amino acids were conserved in the sequences predicted to have bifunctional CPD/(6-4)- activities (Figure 1B).

Considering these results, over a total of 149 sequences we propose that 55 are true homologs of *Sphingomonas* sp. SV9 PhrSph98 (proteins from nodes 1 to 12, Figure 1A), as they conserve all the important amino acids described to be involved in electron transfer, cofactor and DNA lesion binding and stabilization (Figure 1B). A widespread distribution of these enzymes was observed among bacteria, including different genus from proteobacteria (43/55), planctomycete (1/55), bacteroidetes (7/55), acidobacteria (1/55) and cyanobacteria (3/55) (Figure 1A) being the latter the only oxygenic photosynthetic organisms encoding this enzyme.

### 3.2 Phylogeny of CPF proteins

Phylogenetic tree obtained from the CPF proteins was unrooted, with an apparent root on the branch near the group of FeS-BCPs. We manually placed the branches from FeS-BCPs, SPL and bifunctional photolyases as roots in independent trees (Supplementary figure S2). It was found that the simplest tree which has minimum changes across evolution correspond to SPL as root. The maximum likelihood tree reconstruction features 10 main clades. It comprises SPL, CPD PHRs class I, II and III, plant CRY, plant PHR2, ssDNA PHRs, animal CRY and eukaryotic (6–4) photolyases, FeS-BCPs and bifunctional CPD/(6-4)- PHRs (Figure 2C, see supplementary figure S3 for full phylogenetic tree).

**Figure 2.**
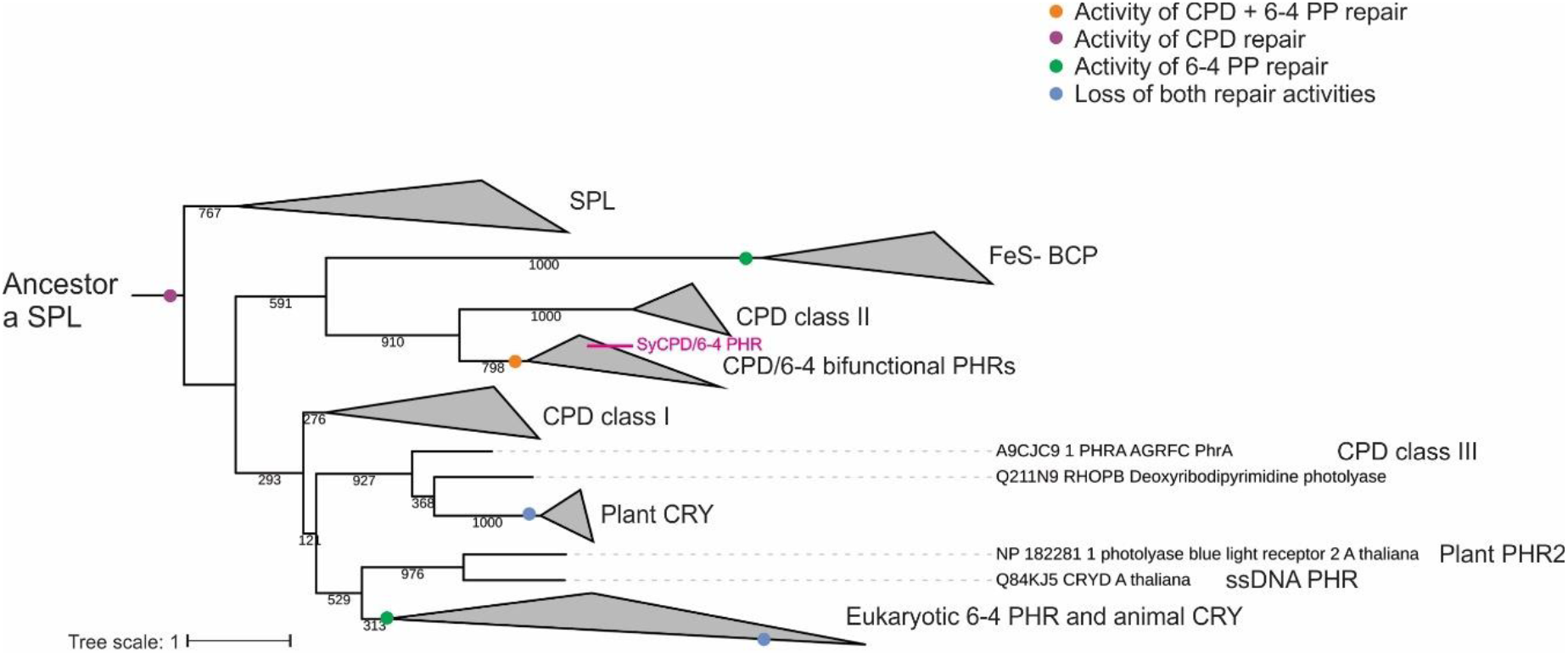
Phylogenetic distribution of the cryptochrome/photolyase family including bifunctional CPD/(6-4)- PHRs. Phylogenetic tree with a CPD photolyase with an iron–sulfur cluster (SPL) as the first common ancestor. Proposed major evolutionary events regarding DNA damage repair activity are indicated by colored dots. Maximum-likelihood probabilities of 1000 replicates are adjacent to each internal node. Node supports below 400 are not shown. Branches corresponding to proteins from the same clade were collapsed. Complete phylogeny is shown in Supplementary Figure S3.

All bifunctional CPD/(6-4)- PHRs constituted one monophyletic group, which was located as a sister group of class II CPD PHRs. This clade was located as a sister group of FeS-BCPs enzymes and all together as sister group of all the rest of the members from the CPF (Figure 2). According to tree inference, FeS-BCPs and eukaryotic 6-4 photolyases lost the CPD activity and gained 6-4 photorepair activity through two independent functional change events, whereas bifunctional CPD/(6-4)- PHRs gained 6-4 activity but also retained CPD repair activity (Figure 2). Results suggest that bifunctional CPD/(6-4)- PHRs are related to class II CPD PHRs, both sharing a common ancestor with FeS-BCPs, in a clade that includes only prokaryotic organisms.

### 3.3 Sequence homology of PhrSph98 and cyanobacteria proteins: structure modeling of bifunctional CPD/(6-4)- photolyase from the cyanobacteria *Synechococcus* sp. PCC 7335

Phylogenetic analysis revealed that canonical CPD/(6-4)- photolyases are also present in a few strains of oxygenic photosynthetic organisms. Blast analysis using PhrSph98 as query towards the *Viridiplantae* group gave no results, indicating that the only oxygenic photosynthetic organisms encoding a bifunctional photolyase were the cyanobacteria *Phormidesmis sp LEGE 1147, Chrondrocistis* sp NIES 4102 and *Synechococcus* sp. PCC 7335 (Figure 1A). Amino acids involved in DNA repair, lesion and cofactor binding were conserved among PhrSph98 and the cyanobacterial homologs (Figure 3A).

**Figure 3.**
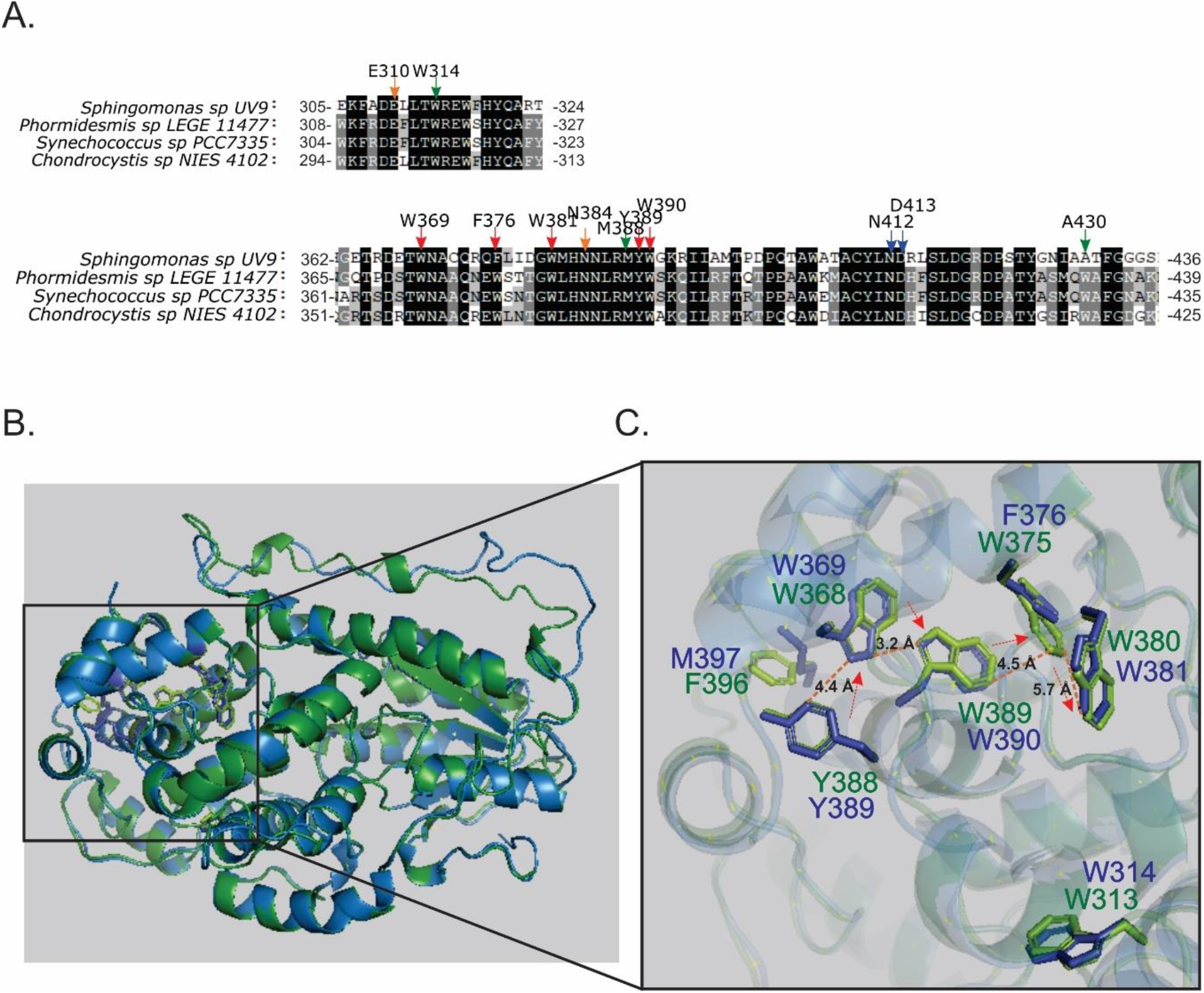
Multiple sequence alignment and molecular modeling. **A.** MSA was performed using MAFFT software. Arrows indicate conserved amino acids involved in electron transfer pathway (red), FAD cofactor binding (blue), DNA lesion binding (green) and lesion stabilization (orange). Conserved residues common to all sequences are shadowed in black and less identity is shown in gray scale. Full MSA is presented in Supplementary Figure S1. **B.** Superposition of predicted crystal tertiary structures from PhrSph98 (blue) and *Synechococcus* sp. PCC 7335 CPD/(6-4)- bifunctional photolyase (green). **C.** Amplification of catalytic domain. Amino acids described that may be involved in electron pathway in PhrSph98 are shown in blue Y389-W369-W390-F376-W381. Homologs from *Synechococcus* sp. PCC 7335 bifunctional CPD/(6-4)- PHR are shown in green. The arrows indicate the electron flux direction. Dash lines indicates distances in amstrongs (Å) among atoms. Figures B and C were generated by using PyMOL.

*Synechococcus* sp. PCC 7335 has several adaptations to different ambient conditions as chromatic acclimation (CA), far-red light photoacclimation (FARLIP) and the presence of a non-canonical nitric oxide synthase enzyme (SyNOS) (Ho et al., 2017; Correa-Aragunde et al., 2018; Herrera-Salgado et al., 2018). In this work, we were particularly interested in the strategies of *Synechococcus* sp. PCC 7335 to cope with UV-B radiation. To analyze that, tertiary structure of PhrSph98 homolog from *Synechococcus* sp. PCC 7335 (herein named as SyCPD/(6-4)- photolyase) was predicted using as template a class II CPD-photolyase from the archaea *M. mazei* (PDB ID: 2XRY) using HHPred online available server. Results obtained show that besides a low percentage of identity of amino acid sequences between PhrSph98 and SyCPD/(6-4)- PHR (40.71%), they adopted similar tertiary structure (Figure 3B). Molecular modeling shown in Figure 3B indicate that the positioning of the amino acids Y389-W369-W390-F376-W381, suggested by Marizcurrena et al. (2020) as important for electron transfer and catalytic activity, were conserved in SyCPD/(6-4)- photolyase. Thus, we propose that SyCPD/(6-4)- PHR may be a true bifunctional CPD/(6-4)- PHR.

### 3.4 The bifunctional CPD/(6-4)- PHR gene in *Synechococcus sp*. PCC 7335 is part of an UV-inducible operon

Genomic context analysis revealed that SyCPD/(6-4) photolyase constitutes a two gene operon assembly with other PHR (herein refer as PHRs operon). This operon is only conserved in *Phormidesmis* sp. LEGE 11477 (accession JADEXO010000055.1, 81.68% of identity and 95% of query coverage) not in *Chrondrocistis* sp NIES 4102. This is the first report of a bifunctional CPD/(6-4) photolyase, as well as a putative PHRs operon in an oxygenic photosynthetic organism.

PHRs operon is encoded in the DNA negative strand (coding sequence from position 38587 to 35587). Two putative −10 (TGCTATACA) and −35 (TTGAAG) boxes were found in the promoter region using the bacterial promoter prediction software BPROM. The analysis of potential transcription factors binding sites revealed the sequences TCACAATT from cyclic AMP receptor protein (CRP) and ACAGACAA from integration host factor (IHF) (Figure 4A). The operon consists of two unidirectional structural genes: an upstream deoxyribodipyrimidine photo-lyase (PHR, WP_038015784.1) and a downstream hypothetical protein (SyCPD/(6-4)- PHR, homolog to PhrSph98, WP_038015781.1) with different translation frames (Figure 4A). Both ORFs overlap in 11 nt within the sequence **ATG**TCAGT**TGA** where TGA is the stop for the upstream gene, and ATG is the start of the second (Figure 4A). We also found a Shine-Dalgarno (SD) intragenic sequence (AGGAG) preceding in seven nucleotides the ATG from the downstream gene. This sequence is absent in the leading gene (Figure 4A). Details of PHRs operon nucleotide sequence is shown in Supplementary figure S4.

**Figure 4.**
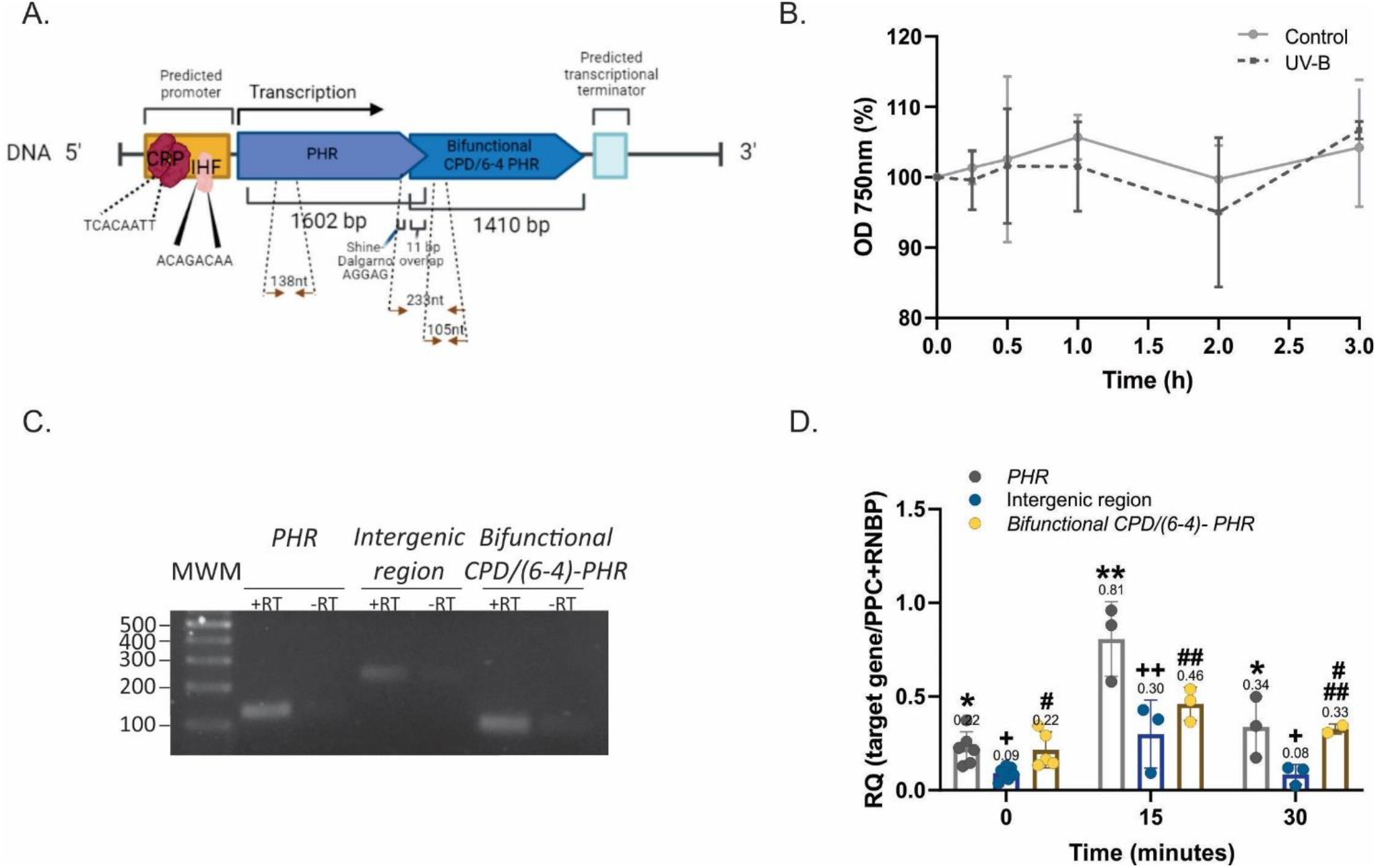
PHRs operon from *Synechococcus* sp. PCC 7335 organization and expression. **A.** Gene organization scheme of the PHRs operon. Drawing was made using BioRENDER tool (https://biorender.com/). Arrows indicate the RT-qPCR regions amplified by primers. Nucleotide sequence from the PHRs operon in shown in Supplementary figure S4. **B.** Transcript characterization. Amplicons from PHR, bifunctional CPD/(6-4)- PHR and the intergenic region (+RT) and negative controls without reverse transcriptase (-RT) were loaded on agarose 2% and subjected to electrophoresis. The gel was stained using 1X Sybr Safe. MWM: molecular weight markers. **C.** Growth curves of *Synechococcus* sp. PCC 7335 strain under control or UV-B treatment for 3h recorded as optical density (OD) at 750 nm. **D.** PHRs expression under UV-B treatment. PHR (grey), SyCPD/(6-4)- PHR (yellow) and the intergenic region (blue) transcript levels of control and UV-B exposed cultures were evaluated by RT-qPCR. Results were expressed as 2^-(ΔCt)^ using as normalizer the geometric mean of PPC and RNBP genes. Normalization of gene expression to individual reference genes in shown in Supplementary figure S5. The One-way ANOVA test was used to determine differences for PHR (*), the intergenic region (+) and bifunctional CPD/(6-4)-photolyase (#) transcript levels at different time treatments (p <0.05). Equal symbols represent no statistical differences.

Photolyases transcript accumulation is induced upon UV-B exposition in several organisms, protecting them from cell death (An et al., 2021; Fernández et al., 2021, 2022; He et al., 2021). First, we analyzed the effect of UV-B (1.6 W m^-2^) on *Synechococcus* sp. PCC 7335 cell culture measuring the OD at 750 nm. As shown in Figure 4B, UV-B do not affect *Synechococcus* sp. PCC 7335 optical density at 750 nm up to 3 h-treatment.

To investigate whether the two PHRs ORFs form a polycistronic mRNA, the intergenic region between PHR and SyCPD/(6-4)- PHR was amplified by qPCR from non-irradiated culture cDNA. Figure 4C shows the amplification of the expected DNA fragment (233nt), confirming the expression of both PHRs as a polycistronic mRNA. The expression of PHR, SyCPD/(6-4)- PHR and the intergenic region were also evaluated during UV-B treatment. Results show that PHR, bifunctional CPD/(6-4)- PHR and the intergenic region expression increased 3.5, 2 and 3-fold respectively after 15 minutes of UV-B exposition. *PHR* transcript was statistically significant higher compared to bifunctional photolyase. After 30 min of UV-B exposure, abundance of all amplicons decreased to control levels (Figure 4D).

## 4 Discussion

### 4.1 Bifunctional CPD/(6-4)- PHRs domain structure is conserved among heterotrophic bacteria and cyanobacteria

Photolyases repair UV-induced DNA damages of CPDs and 6-4 PPs using blue/ UV-A light. The FAD catalytic cofactor, conserved in the whole protein superfamily of photolyase/cryptochromes, adopts a unique folded configuration at the active site and plays a critical role in DNA repair. FADH-functions as the active state to allow efficient electron injection into DNA damage in a process that involves intramolecular electron transfer (Zhang et al., 2017). Based on the *in silico* modeling of PhrSph98 from *Sphingomonas* sp. UV9, Marizcurrena et al., (2020) suggested that electron transfer to FAD involves Y389-W369-W390-F376-W381 amino acids pathway. In class II CPD photolyases, an Asn residue (N403) is highly conserved and involved in FAD cofactor binding. Kiontke et al., (2011) showed that the class II CPD photolyase from *M. mazei* owing the N403A and N403L mutations, have impaired FAD binding. In contrast, the N403D mutant has a FAD incorporation of at least 70% (Kiontke et al., 2011). Also, in class II CPD PHRs, a charge compensation during FADH-photoreduction is performed by H-bonding of the side chain from N403 with D404, whereas in class I this role is taken up by a Glu residue (Kiontke et al., 2011). We showed that all these residues are conserved in PhrSph98 homologs, indicating that bifunctional CPD/(6-4)- PHRs are present in heterotrophic bacteria like proteobacteria, planctomycete, bacteroidete, acidobacteria and in oxygenic photosynthetic bacteria as cyanobacteria. Conservation of class II CPD PHRs residues suggest that these enzymes belong to this class, which was also supported by phylogenetic analysis, positioning bifunctional photolyases as a sister group of CPD class II photolyases.

Additionally, *in silico* tertiary structure model comparison between PhrSph98 and SyCPD/(6-4)- PHR shows that structural arrangement of residues involved in electron transfer to FAD, FAD binding and DNA damages binding are conserved. The tertiary structure conservation of PhrSph98 hallmarks, allows us to suggest that SyCPD/(6-4)- photolyase is a true homolog of this enzyme.

### 4.2 Bifunctional CPD/(6-4)- PHRs clade is a sister group of class II CPD PHRs

Several ancestors for the CPF family have been proposed (Mei and Dvornyk, 2015; Miles et al., 2020; Vechtomova et al., 2020). After analyzing three different phylogenetic tree topologies, we found that positioning of SPL proteins as root retrieves the tree with lesser functional changes during evolution. Accordingly, Xu et al., (2021) suggested that the first common ancestor of the CPF might be a SPL protein, with CPD photolyase activity and a Fe-S cluster, characteristic of ancient proteins. Phylogenetic tree shows that bifunctional CPD/(6-4)- photolyases are a sister group of CPD class II photolyases and both constitute a clade with FeS-BCP proteins, represented only by prokaryotic organisms. 6-4 PP photo repair appear independently in FeS-BCP and bifunctional proteins, probably by mutational events, and conservation of CPD activity is only retained in bifunctional enzymes. A 6-4 PP repair enzyme can be converted into a CPD repair one with only three mutations. However, eleven mutations are needed for the vice versa conversion (Yamada et al., 2016). This asymmetric functional conversion was suggested as a more complex repair mechanism for 6-4 PPs repair (Yamada et al., 2016). Although it seems that conversion of CPD to 6-4 photolyase activity is more difficult in terms of mutations amount, phylogenetic tree supports the origin of the CPF with a CPD repair family.

### 4.3 Characterization of *Synechococcus* sp. PCC 7335 photolyase operon

An operon is a cluster of neighboring genes that are transcribed together and therefore encodes several proteins. Its evolution is hypothesized to be adaptive and towards coordinated optimization of functions (Memon et al., 2013). Results presented here indicate that only the cyanobacteria *Synechococcus* sp. PCC 7335 and *Phormidesmis* sp. LEGE 11477 encode a PHRs operon. It has recently been proposed a new classification of the genus Synechococcus, based in habitat distribution patterns (seawater, freshwater, brackish and thermal environments) that reflects the ecological and evolutionary relationships of its members. According to Salazar et al., (2020), *Synechococcus* sp. PCC 7335 genome did not cluster with any other Synechococcus genome, and match with the Phormidesmiales order, being named *Phormidesmis mexicanus PCC 7335*. This new genus assignation is supported by conservation of PHRs operon both in *Phormidesmis* sp. LEGE 11477 and *Synechococcus* sp. PCC 7335. Thus, the *Phormidesmis* genus is apparently the unique containing a PHRs operon including a bifunctional CPD/(6-4)- PHR reported. Currently, there is a low number of publicly available cyanobacteria genomic sequences (0.6% compared to the total number of genomes available for bacteria and archaea) (Alvarenga et al., 2017). Future genome sequencing of this phylum will allow to determine if this operon occurs in other cyanobacteria.

Operon evolution analysis in cyanobacteria shows that genes in highly and moderately conserved operons code for key cellular processes, as photosynthesis. Contrary, genes in poorly conserved operons may code for functions possibly linked to niche adaptation. Also, newly acquired operons are greater in number, smaller in size, with wider intergenic spacing and weakly coregulated compared to ancient operons. However, a sub-clade comprising the genera *Synechocystis*, *Microcystis*, *Cyanothece* and *Synechococcus* sp. PCC 7002 forms an exceptional case with small intergenic spaces (Memon et al., 2013). *Synechococcus* sp. PCC 7335 PHRs operon matched the criteria of poorly conserved and no intergenic space, indicating that may be a newly formed operon with an adaptative role to protect this organism from high environmental UV-B levels. Here, we observed that UV-B induces the expression of the PHRs operon. This arrangement guarantee that production of the proteins related to DNA repair under UV-B induced damage is simultaneously switched on and off and provides the ability to fast acclimate to new growth conditions. Moreover, there is a potential binding site of the transcription factor CRP upstream of the PHRs operon promoter −35 and −10 boxes. Class I CRP-dependent promoters have this sequence upstream of the promoter boxes. This transcriptional activation involves the interaction between CRP and the carboxy-terminal domain of RNA-polymerase (RNA-P) α-subunit, facilitating the binding of RNA-P to the promoter (Soberón-Chávez et al., 2017). Additionally, a *CRP* knock out strain of *Deinococcus radiodurans* is sensitive to UV radiation, suggesting an important role of CRP in UV protection (Yang et al., 2016). Also, in *D. radiodurans*, a CRP homolog regulates the expression of different DNA repair proteins as *recN* (in response to double-stranded DNA breaks), *pprA* (RecA-independent, DNA repair-related protein) and *uvsE* (UV damage endonuclease that is involved in nucleotide excision repair). Based on this, CRP transcription factor is an interesting candidate to be explored for PHRs operon UV-B inducible expression regulation.

Until now, only two reports describe photolyases encoded in operons. Osburne et al., (2010) characterized an operon of two DNA repair genes (nudix hydrolase-photolyase operon) in the UV hyper-resistant strain marine cyanobacterium *Prochlorococcus* MED4 which has constitutively upregulated expression. Also, Sancarsg et al. (1984), reported an operon of a photolyase with a protein of unknown function in *E. coli*. To our knowledge, our study constitutes the first report that describes an UV-B inducible operon encoding two photolyases, being one of them with a putative bifunctional role in the repair of CPDs and 6-4 PPs damages. We found that upon 15 minutes of UV-B exposition, PHR transcript accumulates faster than bifunctional CPD/(6-4)- PHR. The increase in transcripts level during UV-B irradiation, suggest a role in DNA repairing. The distinct expression of both intra-operonic genes, constitutes a deviation from the generally expected co-expression behavior. A possible explanation may be the occurrence of translational coupling where the juxtaposition of translational stop codon of an upstream gene with the translational start codon of the downstream one reduces the expression levels of the latter and affects mRNA stability (Dogra et al., 2015; Wright et al., 2021). In this sense, the upstream translating ribosome destabilize RNA secondary structure which would prevent optimal expression of the downstream gene (Schümperli et al., 1982). We also found an intragenic SD sequence preceding the bifunctional CPD/(6-4)- photolyase gene, that may be involved in termination-reinitiation of translation. In overlapping genes, the start codon of the downstream gene is typically preceded by a SD motif in most archaea and bacteria (Huber et al., 2019). Cyanobacteria make ample use of the SD motif for reinitiation at overlapping gene pairs while less than 20% of SD sequences are found in the leading gene from operon (Huber et al., 2019). In agreement, no SD sequence was found in the leading gene of PHRs operon.

*Synechococcus* sp. PCC 7335 was isolated from snail shell, from the intertidal zone in Puerto Peñasco, Mexico. As previously mentioned, they have several specific adaptations to prevailing light. For example, they contain chlorophyll *f* that absorb far-red light and go under FaRLiP response and also perform complementary chromatic acclimation. They also contain an operon of PHRs that may contribute to UV-B tolerance, and one of this PHRs correspond to a bifunctional CPD/(6-4)- PHR. Genome analysis of *Synechococcus* sp. PCC 7335 denotes the presence of two other CPF: a cryptochrome/ photolyase family and a deoxyribodipyrimidine photo-lyase (WP_006457129.1, locus 27315-28829 and WP_006453873.1, locus 88140-86491 respectively). BlastP analysis of these two proteins show similarity with cyanobacteria proteins. However, PHRs operon share more similarity with proteobacteria photolyases. In this sense, it is possible that *Synechococcus* sp. PCC 7335 acquire the PHRs operon gene through horizontal gene transfer from proteobacteria. Whether these photolyases have redundant roles or contribute to classify *Synechococcus* sp. PCC 7335 as an UV-B resistant bacteria needs to be explored. Additionally, whether PHRs operon confers an advantage upon UV-B radiation and if CPD/(6-4)- bifunctional photolyase is under horizontally gene transfer are also open questions. Furthermore, it will be interesting to analyze whether repair of two DNA damages by a single protein may confer to *Synechococcus* sp. PCC 7335 an advantage over expressing a CPD and 6-4 PP photolyases. If so, the biochemical characterization of this enzyme may provide useful biotechnological information for the cosmetic and healthcare industries, as photolyases are not encoded by placental mammals and cyanobacteria are emerging organisms as source of bioactive compounds.

## 6 Author Contributions

FMB conceived the original idea. FMB carried out the phylogenies and experiments. LL collaborated in qPCR experiments. FMB, LL, NC and RC contributed to the interpretation of the results. FMB wrote the manuscript. NC and RC contributed to the design of the experiments and critically revised the manuscript. RC supervised the project.

## References

Alvarenga, D. O., Fiore, M. F., and Varani, A. M. (2017). A metagenomic approach to cyanobacterial genomics. Frontiers in Microbiology 8, 809. doi: 10.3389/FMICB.2017.00809/BIBTEX.

An, M., Qu, C., Miao, J., and Sha, Z. (2021). Two class II CPD photolyases, PiPhr1 and PiPhr2, with CPD repair activity from the Antarctic diatom Phaeodactylum tricornutum ICE-H. 3 Biotech 2021 11:8 11, 1–9. doi: 10.1007/S13205-021-02927-0.

Banaś, A. K., Zgłobicki, P., Kowalska, E., Bażant, A., Dziga, D., and Strzałka, W. (2020). All You Need Is Light. Photorepair of UV-Induced Pyrimidine Dimers. Genes 2020, Vol. 11, Page 1304 11, 1304. doi: 10.3390/GENES11111304.

Correa-Aragunde, N., Foresi, N., del Castello, F., and Lamattina, L. (2018). A singular nitric oxide synthase with a globin domain found in Synechococcus sp. PCC 7335 mobilizes N from arginine to nitrate. Scientific Reports 8. doi: 10.1038/s41598-018-30889-6.

Criscuolo, A., and Gribaldo, S. (2010). BMGE (Block Mapping and Gathering with Entropy): A new software for selection of phylogenetic informative regions from multiple sequence alignments. BMC Evolutionary Biology 10, 1–21. doi: 10.1186/1471-2148-10-210/FIGURES/9.

Crooks, G. E., Hon, G., Chandonia, J.-M., and Brenner, S. E. (2004) WebLogo: A Sequence Logo Generator. doi: 10.1101/gr.849004.

Dogra, N., Arya, S., Singh, K., and Kaur, J. (2015). Differential expression of two members of Rv1922-LipD operon in Mycobacterium tuberculosis: Does rv1923 qualify for membership? Pathogens and Disease 73, 29. doi: 10.1093/FEMSPD/FTV029.

Feng, C. Q., Zhang, Z. Y., Zhu, X. J., Lin, Y., Chen, W., Tang, H., et al. (2019). iTerm-PseKNC: a sequence-based tool for predicting bacterial transcriptional terminators. Bioinformatics 35, 1469–1477. doi: 10.1093/BIOINFORMATICS/BTY827.

Fernández, M. B., Latorre, L., Lukaszewicz, G., Lamattina, L., and Cassia, R. (2022). “Nitric oxide-mediated regulation of the physiological and molecular responses induced by Ultraviolet-B (UV-B) radiation in plants,” in Nitric Oxide in Plant Biology doi: 10.1016/b978-0-12-818797-5.00017-0.

Fernández, M. B., Lukaszewicz, G., Lamattina, L., and Cassia, R. (2021). Selection and optimization of reference genes for RT-qPCR normalization: A case study in Solanum lycopersicum exposed to UV-B. Plant Physiology and Biochemistry 160. doi: 10.1016/j.plaphy.2021.01.026.

Fernández, M. B., Tossi, V., Lamattina, L., and Cassia, R. (2016). A comprehensive phylogeny reveals functional conservation of the UV-B photoreceptor UVR8 from green algae to higher plants. Frontiers in Plant Science 7. doi: 10.3389/fpls.2016.01698.

Guindon, S., Dufayard, J.-F., Lefort, V., Anisimova, M., Hordijk, W., Gascuel, O., et al. (2010) New Algorithms and Methods to Estimate Maximum-Likelihood Phylogenies: Assessing the Performance of PhyML 3.0. Available at: http://www.lirmm.fr/~gascuel [Accessed March 6, 2022].

Häder, D. P., Helbling, E. W., Williamson, C. E., and Worrest, R. C. (2011). Effects of UV radiation on aquatic ecosystems and interactions with climate change. Photochemical & Photobiological Sciences 10, 242–260. doi: 10.1039/C0PP90036B.

He, Y., Qu, C., Zhang, L., and Miao, J. (2021). DNA photolyase from Antarctic marine bacterium Rhodococcus sp. NJ-530 can repair DNA damage caused by ultraviolet. 3 Biotech 11, 102. doi: 10.1007/S13205-021-02660-8.

Herrera-Salgado, P., Leyva-Castillo, L. E., Ríos-Castro, E., and Gómez-Lojero, C. (2018). Complementary chromatic and far-red photoacclimations in Synechococcus ATCC 29403 (PCC 7335). I: The phycobilisomes, a proteomic approach. Photosynthesis Research 138. doi: 10.1007/s11120-018-0536-6.

Ho, M. Y., Gan, F., Shen, G., Zhao, C., and Bryant, D. A. (2017). Far-red light photoacclimation (FaRLiP) in Synechococcus sp. PCC 7335: I. Regulation of FaRLiP gene expression. Photosynthesis Research 131. doi: 10.1007/s11120-016-0309-z.

Huang, Y., Niu, B., Gao, Y., Fu, L., and Li, W. (2010). CD-HIT Suite: a web server for clustering and comparing biological sequences. BIOINFORMATICS APPLICATIONS NOTE 26, 680–682. doi: 10.1093/bioinformatics/btq003.

Huber, M., Faure, G., Laass, S., Kolbe, E., Seitz, K., Wehrheim, C., et al. (2019). Translational coupling via termination-reinitiation in archaea and bacteria. Nature Communications 10. doi: 10.1038/s41467-019-11999-9.

Kiontke, S., Geisselbrecht, Y., Pokorny, R., Carell, T., Batschauer, A., and Essen, L. O. (2011). Crystal structures of an archaeal class II DNA photolyase and its complex with UV-damaged duplex DNA. The EMBO Journal 30, 4437. doi: 10.1038/EMBOJ.2011.313.

Kiontke, S., Göbel, T., Brych, A., and Batschauer, A. (2020). DASH-type cryptochromes - Solved and open questions. Biological Chemistry 401, 1487–1493. doi: 10.1515/HSZ-2020-0182/.

Kumar, D., Kannaujiya, V. K., Jaiswal, J., and Sinha, R. P. (2020). Effects of Ultraviolet and Photosynthetically Active Radiation on Phycocyanin of Habitat Specific Cyanobacteria. Journal of scientific research 64. doi: 10.37398/jsr.2020.640110.

Landry, L. G., Stapleton, A. E., Lim, J., Hoffman, P., Hays, J. B., Walbot, V., et al. (1997). An Arabidopsis photolyase mutant is hypersensitive to ultraviolet-B radiation. Proc Natl Acad Sci U S A 94. doi: 10.1073/pnas.94.1.328.

Letunic, I., and Bork, P. (2021). Interactive Tree Of Life (iTOL) v5: an online tool for phylogenetic tree display and annotation. Nucleic Acids Res 49, W293–W296. doi: 10.1093/NAR/GKAB301.

Livak, K. J., and Schmittgen, T. D. (2001). Analysis of Relative Gene Expression Data Using Real-Time Quantitative PCR and the 2-ΔΔCT Method. Methods 25, 402–408. doi: 10.1006/METH.2001.1262.

Marizcurrena, J. J., Acosta, S., Canclini, L., Hernández, P., Vallés, D., Lamparter, T., et al. (2020). A natural occurring bifunctional CPD/(6-4)-photolyase from the Antarctic bacterium Sphingomonas sp. UV9. Applied Microbiology and Biotechnology 104. doi: 10.1007/s00253-020-10734-5.

Mei, Q., and Dvornyk, V. (2015). Evolutionary history of the photolyase/cryptochrome superfamily in eukaryotes. PLoS ONE 10. doi: 10.1371/JOURNAL.PONE.0135940.

Memon, D., Singh, A. K., Pakrasi, H. B., and Wangikar, P. P. (2013). A global analysis of adaptive evolution of operons in cyanobacteria. Antonie van Leeuwenhoek, International Journal of General and Molecular Microbiology 103. doi: 10.1007/s10482-012-9813-0.

Miles, J. A., Davies, T. A., Hayman, R. D., Lorenzen, G., Taylor, J., Mubeena Anjarwalla, ·, et al. (2020). A Case Study of Eukaryogenesis: The Evolution of Photoreception by Photolyase/Cryptochrome Proteins. Journal of Molecular Evolution 88, 662–673. doi: 10.1007/s00239-020-09965-x.

Montgomery, B. L. (2007). Sensing the light: photoreceptive systems and signal transduction in cyanobacteria. Molecular Microbiology 64, 16–27. doi: 10.1111/J.1365-2958.2007.05622.X.

Moon, Y. J., Kim, S. il, and Chung, Y. H. (2012). Sensing and Responding to UV-A in Cyanobacteria. International Journal of Molecular Sciences 13, 16303. doi: 10.3390/IJMS131216303.

Ozturk, N. (2017). Phylogenetic and Functional Classification of the Photolyase/Cryptochrome Family. Photochemistry and Photobiology 93, 104–111. doi: 10.1111/PHP.12676.

Pathak, J., Ahmed, H., Singh, P. R., Singh, S. P., Häder, D. P., and Sinha, R. P. (2019). Mechanisms of Photoprotection in Cyanobacteria. Cyanobacteria: From Basic Science to Applications, 145–171. doi: 10.1016/B978-0-12-814667-5.00007-6.

Pearson, W. R. (2013). An Introduction to Sequence Similarity (“Homology”) Searching. Current protocols in bioinformatics / editoral board, Andreas D. Baxevanis … [et al.] 0 3. doi: 10.1002/0471250953.BI0301S42.

Rastogi, R. P., Sinha, R. P., Moh, S. H., Lee, T. K., Kottuparambil, S., Kim, Y. J., et al. (2014). Ultraviolet radiation and cyanobacteria. Journal of Photochemistry and Photobiology B: Biology 141. doi: 10.1016/j.jphotobiol.2014.09.020.

Rippka, R., Deruelles, J., and Waterbury, J. B. (1979). Generic assignments, strain histories and properties of pure cultures of cyanobacteria. Journal of General Microbiology 111, 1–61. doi: 10.1099/00221287-111-1-1/CITE/REFWORKS.

Scheerer, P., Zhang, F., Kalms, J., von Stetten, D., Krauß, N., Oberpichler, I., et al. (2015). The class III cyclobutane pyrimidine dimer photolyase structure reveals a new antenna chromophore binding site and alternative photoreduction pathways. Journal of Biological Chemistry 290, 11504–11514. doi: 10.1074/JBC.M115.637868.

Schümperli, D., McKenney, K., Sobieski, D. A., and Rosenberg, M. (1982). Translational coupling at an intercistronic boundary of the Escherichia coli galactose operon. Cell 30, 865–871. doi: 10.1016/0092-8674(82)90291-4.

Sinha, R. P., and Häder, D. P. (2008). UV-protectants in cyanobacteria. Plant Science 174. doi: 10.1016/j.plantsci.2007.12.004.

Soberón-Chávez, G., Alcaraz, L. D., Morales, E., Ponce-Soto, G. Y., and Servín-González, L. (2017). The transcriptional regulators of the CRP family regulate different essential bacterial functions and can be inherited vertically and horizontally. Frontiers in Microbiology 8, 959. doi: 10.3389/FMICB.2017.00959/FULL.

Söding, J., Biegert, A., and Lupas, A. N. (2005). The HHpred interactive server for protein homology detection and structure prediction. Nucleic Acids Research 33, W244. doi: 10.1093/NAR/GKI408.

Taboada, B., Estrada, K., Ciria, R., and Merino, E. (2018). Operon-mapper: A web server for precise operon identification in bacterial and archaeal genomes. Bioinformatics 34. doi: 10.1093/bioinformatics/bty496.

Tyagi, R., Srinivas, G., Vyas, D., Kumar, A., and Kumar, H. D. (1992). DIFFERENTIAL EFFECT OF ULTRAVIOLET-B RADIATION ON CERTAIN METABOLIC PROCESSES IN A CHROMATICALLY ADAPTING Nostoc. Photochemistry and Photobiology 55. doi: 10.1111/j.1751-1097.1992.tb04254.x.

Vechtomova, Y. L., Telegina, T. A., and Kritsky, M. S. (2020). Evolution of Proteins of the DNA Photolyase/Cryptochrome Family. Biochemistry (Moscow) 2020 85:1 85, 131–153. doi: 10.1134/S0006297920140072.

Vega, J., Bonomi-Barufi, J., Gómez-Pinchetti, J. L., and Figureueroa, F. L. (2020). Cyanobacteria and Red Macroalgae as Potential Sources of Antioxidants and UV Radiation-Absorbing Compounds for Cosmeceutical Applications. Mar Drugs 18. doi: 10.3390/md18120659.

Wada, N., Sakamoto, T., and Matsugo, S. (2013). Multiple Roles of Photosynthetic and Sunscreen Pigments in Cyanobacteria Focusing on the Oxidative Stress. Metabolites 3, 463. doi: 10.3390/METABO3020463.

Wright, B. W., Molloy, M. P., and Jaschke, P. R. (2021). Overlapping genes in natural and engineered genomes. Nature Reviews Genetics. doi: 10.1038/s41576-021-00417-w.

Xu, L., Chen, S., Wen, B., Shi, H., Chi, C., Liu, C., et al (2021). Identification of a Novel Class of Photolyases as Possible Ancestors of Their Family. doi: 10.1093/molbev/msab191.

Yamada, D., Dokainish, H. M., Iwata, T., Yamamoto, J., Ishikawa, T., Todo, T., et al. (2016). Functional Conversion of CPD and (6-4) Photolyases by Mutation. Biochemistry 55. doi: 10.1021/acs.biochem.6b00361.

Yang, S., Xu, H., Wang, J., Liu, C., Lu, H., Liu, M., et al. (2016). Cyclic AMP Receptor Protein Acts as a Transcription Regulator in Response to Stresses in Deinococcus radiodurans. doi: 10.1371/journal.pone.0155010.

Zhang, M., Wang, L., and Zhong, D. (2017). Photolyase: Dynamics and electron-transfer mechanisms of DNA repair. Archives of Biochemistry and Biophysics 632. doi: 10.1016/j.abb.2017.08.007.

