## Supplementary material for "A bifunctional CPD/(6-4)- photolyase from the cyanobacteria *Synechococcus* sp. PCC 7335 constitutes an UV-B inducible operon: new insights into photolyases phylogenetic evolution"

### Table 1. Sequences and molecular characteristics of the primers used in this study.

| Gene symbol | Accession No. | Official Full Name (MGI) | Primer Pair (5′–3′) | Amplicon length (bp) | Primer efficiency (%) | $R^2$ | Amplicon Tm (°C) | Reference |
|---|---|---|---|---|---|---|---|---|
| *PHR* | WP_038015784.1 | Deoxyribodipyrimidine photo-lyase | Fw GGCAGTATTTGTTCCTCAAGG<br>Rv GCTCGATATTGCCGTATGT | 138 | 90 | 0.99 | 78.22 | This paper |
| *Bifunctional CPD/(6-4)-PHR* | WP_038015781.1 | Hypothetical protein | Fw CGGGTTCGCTCAAAGCAG<br>Rv GCTCGGTTGAGGATCACTA | 105 | 89 | 0.99 | 77.52 | This paper |
| Intergenic region | - | - | Fw AGGCTTCGGCTACTTCC<br>Rv GCTCGGTTGAGGATCACTA | 233 | 88 | 0.99 | 79.72 | This paper |
| *RNPB* | - | RNase P RNA | Fw CGGCTCAAAGCAAGGCTCAA<br>Rv GATGCGATTACGGACGACTGC | 123 | 93 | 0.99 | 79.45 | Correa- Aragunde et al., 2018 |
| *PPC* | WP_198011378.1 | Phosphoenolpyruvate carboxylase | Fw CACCCTGCCCGAATTATCGGTAC<br>Rv CCACGTAACGTCAGGAGTGACAG | 151 | 90 | 0.99 | 76.52 | This paper |

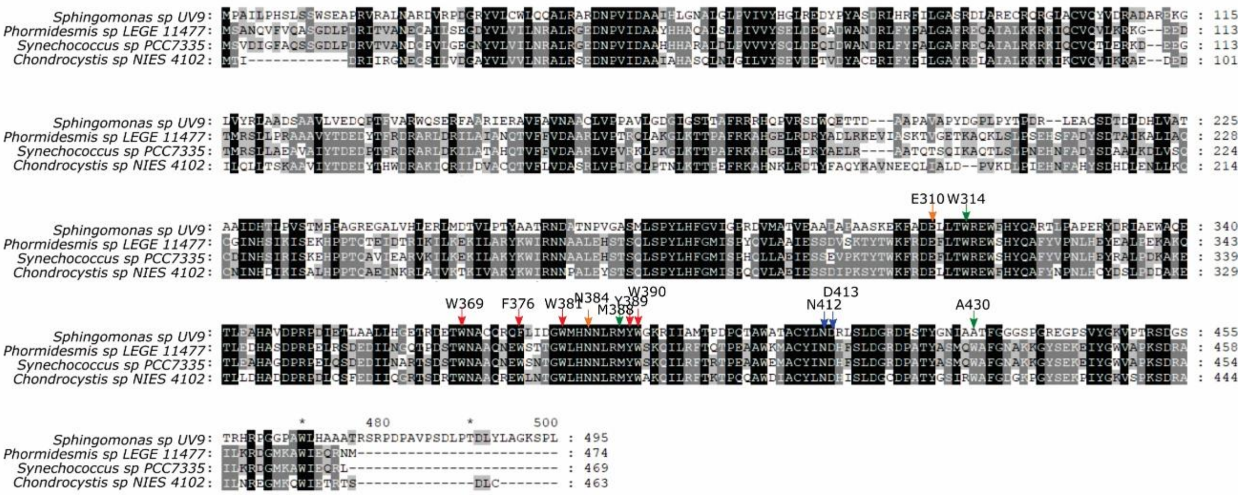


**Figure S1. MSA from *Sphingomonas* SV9 CPD/6-4 bifunctional photolyase homologs in cyanobacteria.** MSA was performed using MAFFT software. Arrows indicate conserved amino acids involved in electron transfer pathway (red), FAD cofactor binding (blue), DNA lesion binding (green) and lesion stabilization (orange). Conserved residues common to all sequences are shadowed in black and less identity is shown in gray scale.

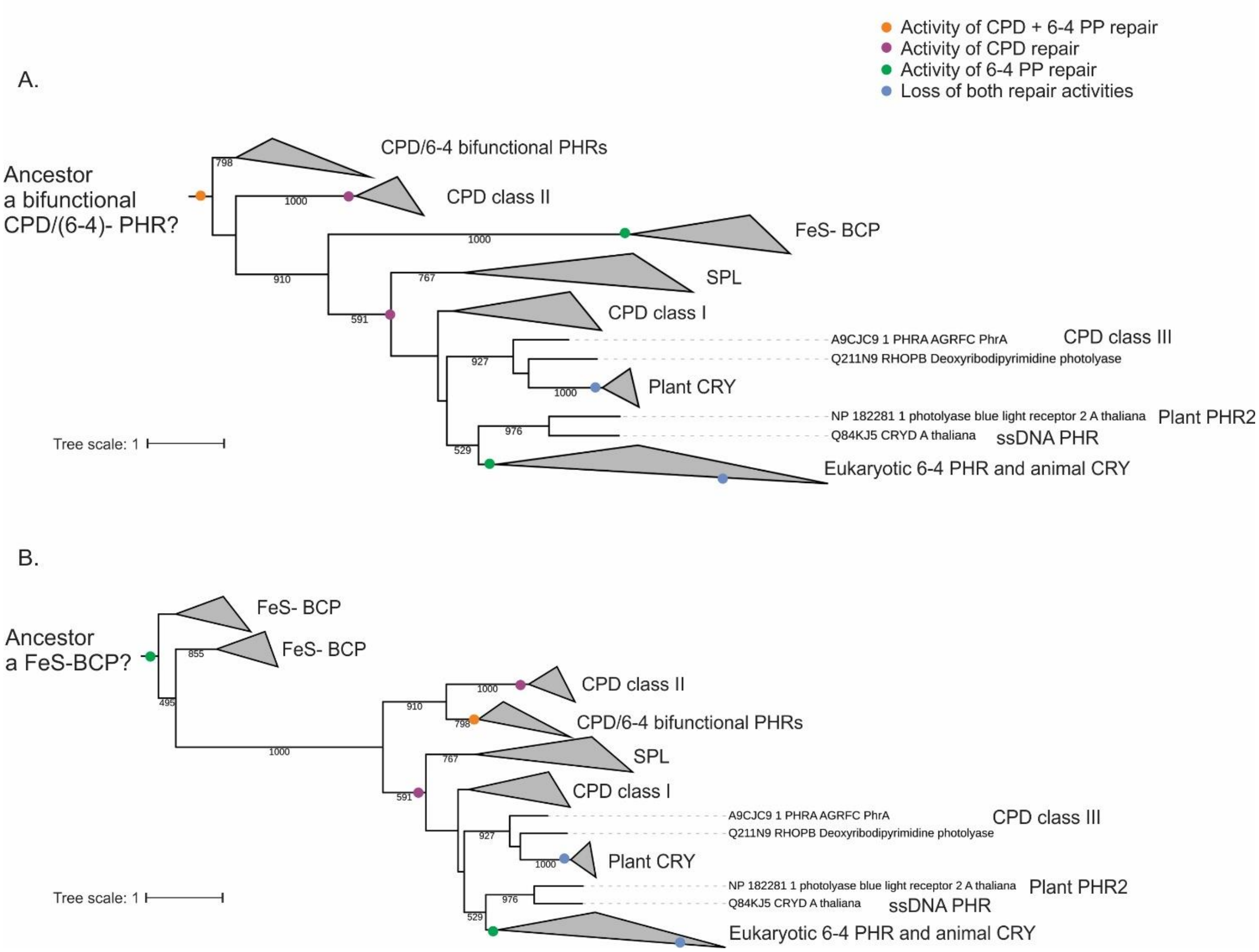


**Figure S2. Other possible evolutionary pathways of CRY/PHR family.** The first common ancestor of CRY/PHR family might be (*A*) bifunctional CPD/(6-4)- PHR, (*B*) a 6-4 photolyase with an iron–sulfur cluster like a FeS-BCP. Proposed major evolutionary events regarding DNA damage repair activity are indicated by colored dots. Maximum-likelihood probabilities of of 1000 replicates are adjacent to each internal node. Node support below 400 are not shown.

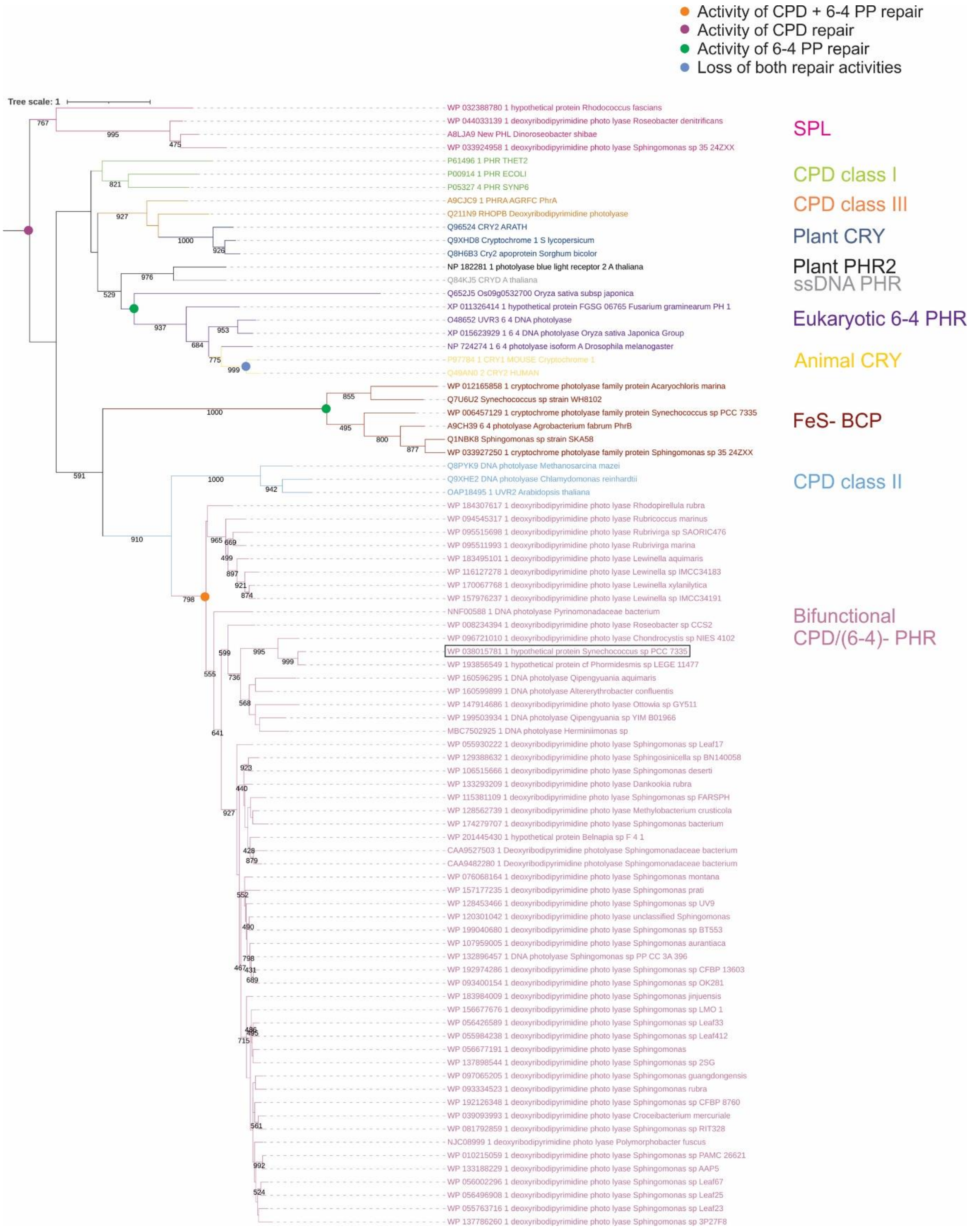


**Figure S3. Possible evolutionary pathways of CRY/PHR family.** Phylogenetic tree with a CPD photolyase with an iron–sulfur cluster (SPL) as the first common ancestor. Proposed major evolutionary events regarding DNA damage repair activity are indicated by colored dots. Maximum-likelihood probabilities of the node support below 400 are not shown. Bootstrap values of 1000 replicates are adjacent to each internal node. Position of SyCPD/(6-4)- PHR is indicated with a rectangle.

**A.**

-35 -10

TTACCAGAACAACAAACACAATCACAATTGCCTTCTTGAAGCGTATGCATCCGCGCTTTG

CTATACAGGCGCTTACAGACAAAAGTAAAAAGTAAGTAGGAAGGATGATGCTCTGGGCAC

CCAGAATCTTCTAATGTTTGAATAAAATAGCGCTGTTTGATTCGAAGGACTTATTTCTTC

TTGATTGACGCTGCTGTTTTGATCAGACCCTTACCGTTAGCGGATAGCAATTGATGAGCG

CTGTACAAGTTGTGTGGTTCAAAAAAGACTTTCGACTCTTCGATCATCGCTGCTTGACGT

TCGCTGCGCAAAAGGGACCGGTTCTACCGCTTTATGTTGTCGAGCCAGACTACTGGGCAC

TAGAAGATACGAGCTATCGGCAGTATTTGTTCCTCAAGGGCTGTGTAGAAGAAACGATTG

AGGCATTAGAAGAGCTGGGTGGGTCGATGGCGATCGCCTCTGGTGATGTAGTAGAGGTGC

TCAGTCGACTCCAGAAAACATACGGCAATATCGAGCTTTGGGCACATCAGGAAACAGGCA

ACAACTGGACGTTTCAACGGGATAAAGCGGTTAGAAAGTGGTGCCAGCAAGCAGGGGTAG

CCTTCCATGAGCCGCTTCAGTTTGGCGTATGGCGAGGCTCCAAAATAAACCGCGATCACT

GGGCTAAGCAGTGGGATTCGCTGATGGCAGAACCCATCGCATCACTGCCAGAAAGCATCT

CTTTTGTAGCGCATTCTGAAGATAGAACGATGCCATCGCCCGAAGTACTAGGGCTGAAGC

ACGATGGCATCACCGCACTGCAGCCACCCGGTAGAAACGCAGCGCTTCAGGTTCTAAGCA

GTTTTTTGTACGAACGCGGCGAGAGATATCGTACAGATATGTCTTCGCCTGTGACTGGTG

AAAGTGGCTGCTCTAGGCTTTCTCCATACCTGGCCTACGGTGCGATCTCCATGCGCGAAA

CCTATCAAGCCACTCAAAAAAGAATTGCCGAAGTCGCTGCTATGCCCAAAGCAGAAAGGG

GGACCTGGGCCGGATCGCTCTCTTCTTTTGTGGGCCGTCTGCACTGGCACTGTCATTTCA

CTCAAAAGCTAGAGCTAGAGCCTGAACTTGAGTGGCTACCCATGGCAAGAGCCTATACAG

GCATCCGAGACGACGGCGACCATGCAATTAAGCTACGAGCGTTTGCAGAAGGTCAAACTG

GCTACCCCTTTGTTGATGCCTGTATGCGTTACCTGCGCGCGACCGGTTGGATTAACTTCC

GGATGCGAGCGATGCTGATGAGCTTTGCTAGCTACGACCTATGGCTGCCCTGGCAAAAGA

GTGGCGATGTTCTAGCCAGGCTTTTCACCGATTATGAGCCTGGCATCCACTGGCCTCAAT

CTCAGATGCAATCAGGCGTCACAGGTATCAACGCTATCCGGATCTATTCACCTATCAAAC

AAGGGCTAGACCAAGATATAGAAGGGACGTTTACCCGCCAGTGGGTTCCAGAACTCGCCG

CACTACCTGACGAAATACTACAGACACCTTGGCTCTTAGAAGGAGAACTCTCGTATCCGC

CGCCCATTGTGGAACATAAAGAGGCTGCAGCCTTAGCAAAGTCTAGGCTCTGGGCAATCA

AGAAAACACCTGAAGCAAAGAAAGAAGCTGAGCAAGTATACGAAAAGCACGGATCTAGAA

AGAAACCTCGTCGCAGAACTTCTTCAAGAAAGGCTTCGGCTACTTCCGCTCAGCGCACAA

AAGCTAAGCGTACTTCAAAAGGCAAGGCAAGCGCACAAAAAACGACTGAAAAACTCCCTA

CAAAAGCTGCTCAGGAGGCTGCTC**ATG**TCAGT**TGA**TATCGGGTTCGCTCAAAGCAGCGGC

GATCTGCCTGATAGAGTGACTGTTGCGAATGATCAACCTGTTTTGGGAGAGGGCAACTAT

GTTCTAGTGATCCTCAACCGAGCCTTACGAGGTGAAGACAACCCGGTCATAGACGCAGCA

CACCATCACGCGCGGGCGCTGGATTTACCTGTTGTTGTTTACAGCCAGTTAGACGAGCAG

ATCGATTGGGCGAATGACCGTCTGTTCTACTTTGCTTTGGGTGCGTTTAGAGAATGTGCG

ATCGCCCTTAGAAAGCGCAAAATCCAGTGCGTACAGACTCTAGAGCGAAAGGATGAGGAA

GGGACTATGCGATCGCTGCTCGCCGAAGCCGTCGCCATCTATACCGACGAAGATCCTACC

TTTCGAGATAGGGCTAGATTAGACAAAATTCTAGCAACTGCCCATCAAACCGTATTCTTT

GTAGACGCCGCTAGACTTGTTCCTGTCCGCAAGCTGCCAAAAGGGCTCAAGACAACCCCT

GCTTTTAGAAAAGCGCACGGAGAGTTGAGAGAGCGTTATGCTGAGCTGCGAGCAGCGACC

CAAACCAGTCAGATCAAGGCACAAACGCTATCGCTACCAAACGAACACAACTTCGCAGAC

TACAGCGACGCAGCTCTCAAAGATCTGGTTTCACAGTGCGATATCAATCACAGTATCAGA

ATCTCGAAAGAGCACCCACCGACTCAAGCAGTCATAGAAGCTCGAGTCAAAATCCTAAAA

GAGAAAATCCTTGCCAAATACAAATGGATTAGAAACAACGCAGCACTAGAACATTCCACC

TCACAGCTATCCCCCTATCTACACTTTGGCATGATTAGCCCGCACCAGCTCCTAGCTGAA

ATTGAATCTTCAGAGGTTCCCAAAACTTACACCTGGAAATTTCGAGACGAATTCCTAACC

TGGCGAGAATGGTCCCACTATCAGGCCTTCTACGTTCCTAATCTGCATGAATACAGGGCC

TTACCAGACAAAGCAAAAGAGACGCTAGAGGCTCATGCAGGCGATCCACGCCCCGAGCTG

CAATCAGATGAAGACATTTTGAACGCTCGCACGTCCGATAGTACCTGGAATGCTGCTCAA

AACGAGTGGTCGAATACGGGATGGCTACACAACAACCTGCGGATGTACTGGTCAAAACAA

ATCTTACGCTTTACCCGAACACCAGAAGCCGCCTGGGAAATGGCTTGCTACATCAATGAC

CATTTCTCTTTAGATGGCAGAGATCCTGCAACCTATGCCTCTATGCAGTGGGCCTTCGGT

AATGCGAAAAAAGGCTACAGCGAGAAGGAGATTTACGGATGGGTAGCACCGAAGTCTGAC

AGAGCTATCTTGAAAAGAGACGGCATGAAAGCATGGATTGAGCAAAGGCTATAGAGAGCC

AAATCCCTTCTTTTTTTTTGTTCTTCTTTTTCTTCTTTCCACCGCCAGCGCCACGCCAACC

GGGCCTAGGCGCCCCACCGAAAGGATTTCTCCCCATCGGTCCGCCACCCATTGGACCGCC

GCCCATACCAGGAAAGCTACCGTCAAATCCTGGTAGCCCCATATTTCCTTGGCCCATTTG

CTGCATGAGTCCGCGCATCCGCTGAAAATCACTCACCAGCTTGCCGACATCCTTCTCGGC

ATAGCCAGCACCCTTTGCCACCCGCCGCCGCCGACTAGGCGAACCGGACAACACTTCTGG
GTTCTGCCGTTCTTCCATCGTCATCGAATTGATCATGGCCTCACAGCGCTTTAGCTGCTG
ATGACGATGGAAGAACGGCAG

**B.**

| Operon | IdGene | Type | COGgene | PosLeft | postRight | Strand | Function |
|---|---|---|---|---|---|---|---|
| 1 | | | | | | | |
| | IFGKMJPH_00001 | CDS | COG0642 | 93 | 4418 | + | [T] Signal transduction histidine kinase |
| 2 | | | | | | | |
| | IFGKMJPH_00002 | CDS | COG0415 | 4674 | 6275 | + | [L] Deoxyribodipyrimidine photolyase |
| | IFGKMJPH_00003 | CDS | COG0415 | 6265 | 7674 | + | [L] Deoxyribodipyrimidine photolyase |

**Figure S4. Genetic organization of the PHRs operon. A.** In blue, the PHR gene and in dark blue the bifunctional CPD/(6-4)- PHR. In pink, the overlapping region between both genes. In grey, the Shine Dalgarno sequence. The ATG from downstream gene and TGA from upstream gene are shown in bold. Underlined, primers used for RT- qPCR analysis. -35 and -10 boxes are indicated at the promoter region. **B.** Operon mapper prediction of prokaryotic operons. IFGKMJPH_00002 corresponds to PHR and IFGKMJPH_00003 to SyCDP/(6-4) predicted bifunctional PHR.

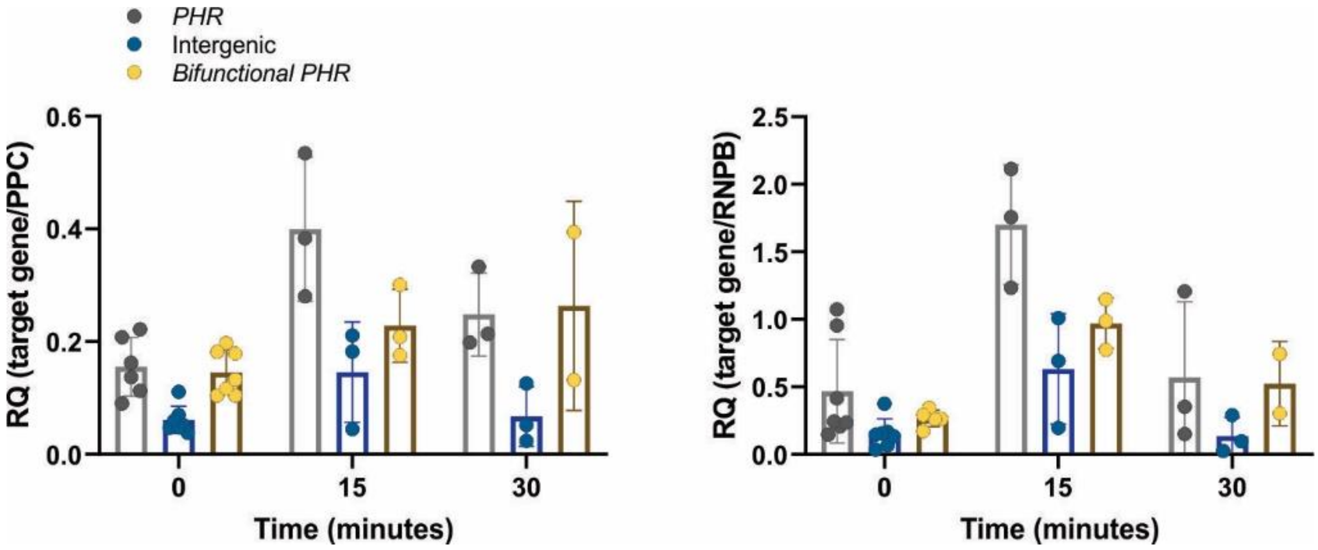


**Figure S5. Operon expression analysis under UV-B treatment by RT- qPCR.** PHR (grey), SyCPD/6-4 PHR (yellow) and the intergenic region (blue) transcript levels under control or UV-B treatment were evaluated by RT-qPCR. Results were expressed as $2^{-(\Delta Ct)}$ using as normalizer PPC (left panel) or RNPB (right panel).
